# Estimates of vital rates and predictions of population dynamics change along a long-term monitoring program

**DOI:** 10.1101/186932

**Authors:** Simone Vincenzi, Dusan Jesensek, Alain J. Crivelli

## Abstract

Despite the widespread recognition of the importance of monitoring, only a few studies have investigated how estimates of vital rates and predictions of population dynamics change with additional data collected along the monitoring program. We investigate how using the same models estimates of survival, individual growth, along with predictions about future population size change with additional years of monitoring and data collected, using as a model system freshwater populations of marble (*Salmo marmoratus*), rainbow (*Oncorhynchus mykiss*), and brown trout (*Salmo trutta* L.) living in Western Slovenian streams. Fish were sampled twice a year between 2004 and 2015. We found that in 3 out of 4 populations, a few years of data (3 or 4 sampling occasions, between 300 and 500 tagged individuals for survival, 100 to 200 for growth) provided the same estimates of average survival and growth as those obtained with data from more than 15 sampling occasions, while the estimation of the range of survival required more sampling occasions (up to 22 for marble trout), with little reduction of uncertainty around the point estimates. Predictions of mean density and variation in density over time did not change with more data collected after the first 5 years (i.e., 10 sampling occasions) and overall were within 10% of the observed mean and variation in density over the whole monitoring program.

## 1. Introduction

The estimation of vital rates and life-history traits and how they vary with habitat and population factors are crucial both for our understanding of population dynamics, risk of extinction, and evolution of traits in natural populations, and for informing management strategies in conservation programs (Frederiksen et al., 2014). To understand how variation in vital rates and life histories of organisms among individuals and through time emerge and how that variation contributes to population dynamics and risk of extinction, we typically need long-term monitoring studies that include contrasting environmental conditions (Elliott, 1994), longitudinal data (Thomson et al., 2009), and statistical models that can tease apart environmental and biological contributions to the observed temporal (and spatial, in case of meta-populations or multiple populations) variation in vital rates, life histories, and population dynamics (Letcher et al., 2015).

Monitoring can be defined as the systematic acquisition of information over time (Gerber et al., 2005). When the goal is informing management strategies for conservation of species, monitoring is the process of collecting information about state variables (e.g., abundance, size, vital rates, life histories) at different points in time and space for the purpose of detecting changes in those variables through time, and among space and individuals (Gerber et al., 2005).

Also in the context of conservation biology, the purpose of a scientific investigation should drive model formulation and the type and amount of data collected. It follows that defining “long-term” for monitoring is often challenging and always context-dependent, since how long the monitoring of natural populations must be carried out depends on the generation time and longevity of the organisms, the characteristics of the environment in which the species lives, and the goals of the monitoring program. For instance, multiple generations are needed to test hypotheses on the evolution of life-history traits or on mechanisms of recovery of species at risk of extinction (Vincenzi et al., 2017). However, as stated by Elliott (1994) in his comments on the value of long-term monitoring in the influential book “Quantitative Ecology and the Brown Trout”, the recovery of a forest ecosystem from clear cutting may last 20 years but this is only about 5% of the time required for the forest to reach steady-state. On the other hand, a study on bacteria may last only a few days, but include many generations. For conservation programs, sufficient knowledge to address most practical problems related to conservation and management of endangered species will usually be obtained within a few years or generations of the monitored species, after which the cost of monitoring begins outweighing the expected benefits with regard to management strategies and overall decision-making (Possingham et al., 2012).

Within a population, habitat factors - both extrinsic (e.g., weather, food) and intrinsic (e.g., population density) - and their interaction (Baerum et al., 2013), determine a large part of the temporal variation in the distribution of vital rates, in recruitment, and in population size, age-, and size-structure (Jonsson and Jonsson, 2011). If the fundamental parameters of an ecological system are constant, i.e. if habitat factors vary little through time and/or space, then monitoring for learning in the context of conservation biology will rarely be long-term (Possingham et al., 2012). On the other hand, highly stochastic environments such those characterized by the occurrence of extreme events (whose “extremeness” is due to a combination of rarity and intensity of the event, Vincenzi 2014) require decades-long monitoring in order to capture them and their effects on vital rates, life histories, population dynamics, and risk of population extinction (Vincenzi et al., 2017). In addition, serendipitous findings and an appreciation of the effects of subtle variation in life histories in natural populations of long-lived species on individual and population processes may only come after many years of monitoring, although one might expect new knowledge to be gained in ever decreasing increments (Possingham et al., 2012). Lastly, especially for small population, many years of data may be necessary to reduce the uncertainty around the estimation of vital rates due to sample size effects (Reynolds, 2012).

Despite the widespread recognition of the importance of monitoring, only a few studies have investigated how estimates of vital rates and predictions of population dynamics change with additional data collected through the monitoring program, and what are the minimum or - when factoring in the costs in money and time of monitoring - optimal years of monitoring or amount of data collected for estimating vital rates and predicting population dynamics (Caughlan and Oakley, 2001). For instance, Gerber et al. (2005) studied how long should we monitor the recovery of an over-fished stock to determine the fraction of that stock to reserve, and they found that the optimal monitoring time frame is rarely more than 5 years. After 5 years, the expected benefit of reduced uncertainty about parameters in the system was negligible compared to the expected gain from earlier exploitation. Other studies have investigated how using the same data and different models of population dynamics (e.g., matrix selection *vs*. element selection) can provide different predictions in the context of Population Viability Analysis (e.g., Bell et al. 2013).

In this work, we investigate how, when using the same models, estimates of survival and body growth, along with predictions about future population size, change with additional years of data from monitoring. We use as a model system freshwater populations of marble (*Salmo marmoratus*), rainbow (*Oncorhynchus mykiss*), and brown trout (*Salmo trutta* L.) living in Western Slovenian streams. These trout populations have been monitored (tag-recapture) since 2004 as part of the ongoing conservation program for the endangered marble trout (Crivelli et al., 2000). The conservation program had the goals of genetically rehabilitating marble trout through the eradication of alien salmonids, acquiring information on variation in vital rates and life histories of the species, and predicting variation in risk of population extinction among the eight genetically pure marble trout populations living in the Western Slovenia (Crivelli et al., 2000; Vincenzi et al., 2016b). In particular, estimates of survival and body growth are fundamental to any assessment of population demographics and population dynamics for management.

We estimated average and time-specific survival probabilities and average growth trajectories for each year of sampling, i.e. with cumulative tag-recapture data up to 2006, 2007, and so on up to 2014, and then used models of population dynamics to study how predictions of mean population size and its temporal variation change with additional years of sampling data. Due to the similarity of the monitored species (all belonging to the family *Salmonidae*) and their restricted geography (Western Slovenia), our results are more descriptive than prescriptive. We encourage the undertaking of similar analyses by other conservation scientists and practitioners, with the goal of providing general guidelines on the minimum length of monitoring programs, amount of data collected, or individuals tagged and sampled for goals ranging from the estimation of vital rates to prediction of population dynamics and extinction risk.

## 2. Material and Methods

### 2.1. Study area and species description

We estimated survival probabilities and growth trajectories, and predicted population dynamics for the marble trout populations of Lower Idrijca [LIdri_MT] and Upper Idrijca [UIdri_MT] (Vincenzi et al., 2016b), rainbow trout population of Lower Idrijca [LIdri_RT] (Vincenzi et al., 2017a), and brown trout population of Upper Volaja [UVol_BT] (Vincenzi et al., 2017b). In LIdri, marble trout [LIdri_MT] live in sympatry with rainbow trout [LIdri_RT] (Vincenzi et al., 2017a; Vincenzi et al., 2011). Both UIdri_MT and UVol_BT live in allopatry. LIdri_RT was created in the 1960s (Vincenzi et al., 2017a) and UVol_BT in the 1920s (Vincenzi et al., 2017b) by stocking rainbow and brown trout, respectively. Both populations have been self-sustaining since their creation (Vincenzi et al., 2017a, 2017b).

### 2.2. Sampling

Populations were sampled bi-annually in June and September. The first sampling for LIdri_MT, LIdri_RT, and UIdri_MT was in June 2004, and in September 2004 for UVol_BT. Sampling protocols are described in greater details in Vincenzi et al. (2016b) and Vincenzi et al. (2017b). If captured fish had *L* > 115 mm, and had not been previously tagged or had lost a previously applied tag, they received a Carlin tag (Carlin, 1955) and age was determined by reading scales. Fish are aged as 0+ (juveniles) in the first calendar year of life, 1+ in the second year and so on. Sub-yearlings of marble, rainbow, and brown trout are smaller than 115 mm in June and September, so fish were tagged when at least aged 1+. The adipose fin was also removed from all fish captured for the first time (starting at age 0+ in September), including those not tagged due to small size at age 1+. Therefore, fish with intact adipose fin were not sampled at previous sampling occasions at age 0+ or 1+.

We estimated density of fish older than 0+ using a two-pass removal protocol (Carle and Strub, 1978) as implemented in the R (R Development Core Team, 2014) package FSA (Ogle, 2015). Total stream surface of the monitored area (1084 m^2^, 1663 m^2^, and 746.27 m^2^ for LIdri, UIdri, and UVol, respectively) was used for the estimation of fish density (in fish ha^−1^).

### 2.3. Statistical analysis of survival and growth

Our goal was to investigate how estimates of (a) average and time-specific survival probabilities and (b) average body growth, and (c) predictions of population dynamics change with each additional year of sampling data, where *Y*_f_ is the last year of monitoring/data collection in September. As simulations of population dynamics often prevent the use of null-hypothesis testing and multiple comparisons increase the “researcher degrees of freedom”, including the choice of convenient hypotheses to test (Gelman and Loken, 2013), we present and discuss our results on variation in survival, growth, and population dynamics from a qualitative point from a view, that is without formal null-hypothesis testing.

For each population, the first models were estimated with *Y*_f_ = 2005, i.e. using data up to from September 2005. For the analysis of survival, we used both June and September data, while for the analysis of growth we used only September data. The cumulative sum of tagged fish in September of each year increased approximately linearly through time in all four populations (Fig. 1).

**Fig. 1.**
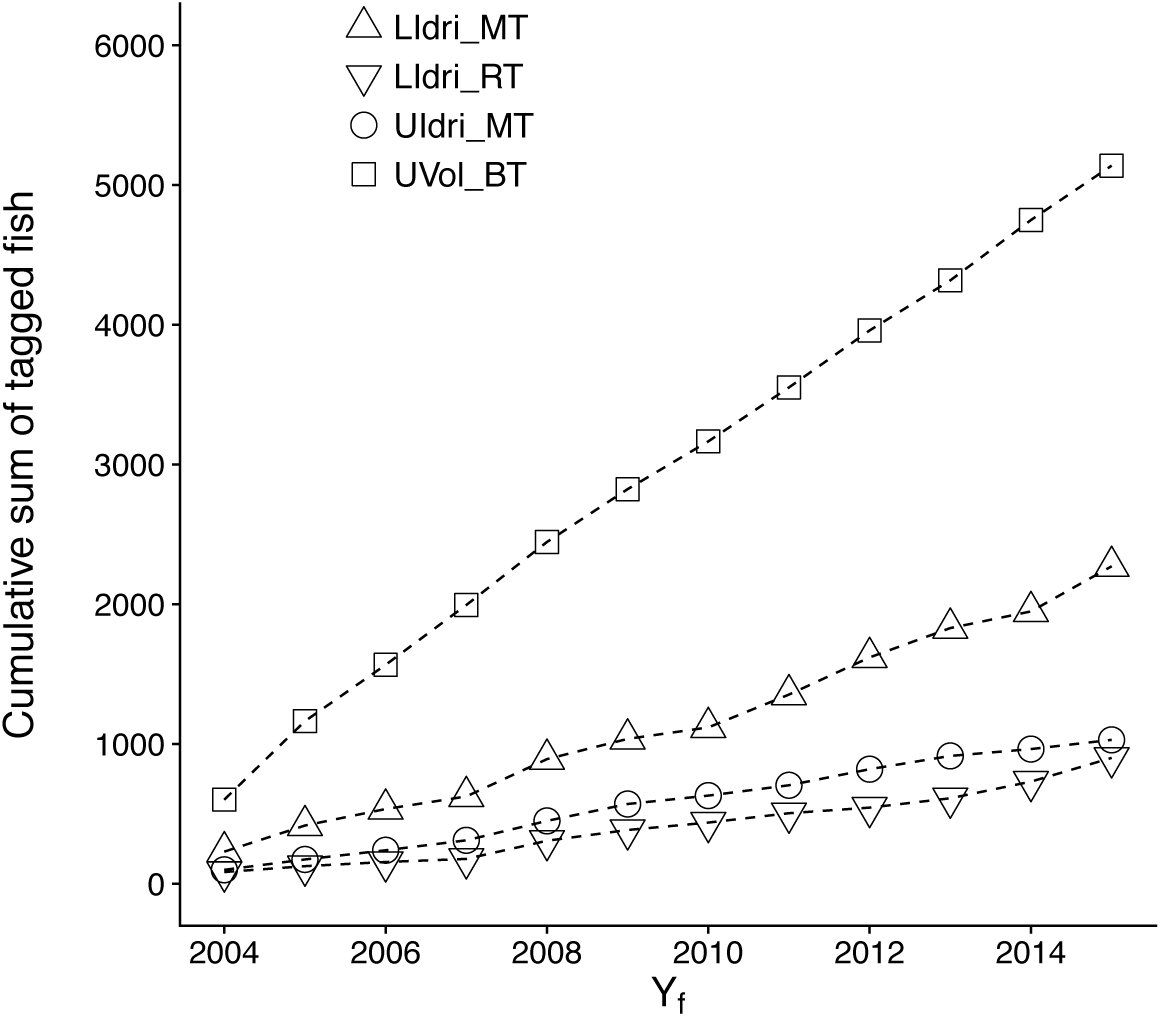
Cumulative sum of tagged fish along the monitoring program from 2004 to 2015. Fish were tagged when at least 115-mm long.

#### Survival

Two relevant probabilities can be estimated from a capture history matrix: *ϕ*, the probability of apparent survival, and *p*, the probability that an individual is captured given that it is alive (Thomson et al., 2009). We used the Cormack–Jolly–Seber (CJS) model as a starting point for the analyses (Thomson et al., 2009). We tested the goodness-of-fit of the CJS model with the program Release. Previous work has found no or minor effects of population density, water temperature, body size, or sex on survival in marble trout (Vincenzi et al., 2016b). Other studies on the same marble (Vincenzi et al., 2016b), rainbow (Vincenzi et al., 2017a), and brown trout populations (Vincenzi et al., 2017b) have found average probability of capture of tagged fish to be > 0.6.

We modeled the seasonal effect (*Season*) as a simplification of full time variation, by dividing the year into two periods: June to September (*Summer*), and the time period between September and June (*Winter*). Since length of the two intervals (*Summer* and *Winter*) was different (3 months and 9 months), we estimated probability of apparent survival on a common annual scale.

In order to compare model results when different data were used, models tested included for either only the constant term (i.e., average apparent survival over all the sampling intervals) or sampling occasion. For probability of capture *p*, following Vincenzi et al. (2016b) we tested models with either *Age*, *Season*, *Cohort*, or sampling occasion as predictors, along with the capture model with no predictors (i.e., constant probability of capture).

We used Akaike Information Criterion (AIC) for model selection (Akaike, 1974). For each *Y*_f_, we extracted average survival probabilities over the whole sampling period (2004 to *Y*_f_) and for each sampling interval from the respective best models of (a) average over the whole sampling period 2004 to *Y*_f_, and (b) for each sampling interval from 2004 to *Y*_f_. We carried out the analysis of survival using the package *marked* (Laake et al., 2013) for R (R Development Core Team, 2014).

#### Growth

The standard von Bertalanffy Growth Function (vBGF; von Bertalanffy 1957) is

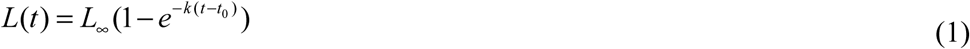

where *L*_∞_ is the asymptotic size, *k* is a coefficient of growth (in time^−1^), and *t*_0_ is the (hypothetical) age at which length is equal to 0.

In the vast majority of applications of the vBGF, *L*_∞_, *k*, and *t*_0_ have been estimated at the population level starting from cross-sectional data, without accounting for individual heterogeneity in growth due to genetic, environmental, and stochastic factors. However, when data include measurements on individuals that have been sampled multiple times, failing to account for individual variation in growth may lead to biased estimations of asymptotic size and mean length-at-age (Vincenzi et al., 2016a, 2014).

In this study, we used the formulation of the vBGF specific for longitudinal data of Vincenzi et al. (2014b), in which *L*_∞_ and *k* may be allowed to be a function of shared predictors and individual random effects. However, in this study we limited our analyses to models including only the intercept (i.e., the overall mean) and individual random effects, i.e. we did not include group effects. In the estimation procedure, we used a log-link function for *k* and *L*_∞_, since both parameters must be non-negative. We set:

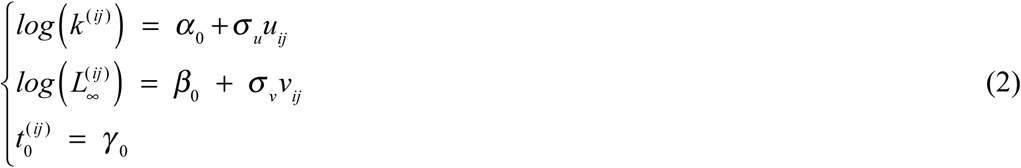

where *u* ~ *N*(0,1) and *v* ~ *N*(0,1) are the standardized individual random effects, *σ*_*u*_ and *σ*_*v*_ are the standard deviations of the statistical distributions of the random effects, *i* is the individual. Since the growth model operates on an annual time scale and more data on tagged fish were generally available in September of each year, we used September data for modeling lifetime growth.

Models were fitted with the Automatic Differentiation Model Builder (ADMB), an open source statistical software package for fitting nonlinear statistical models (Bolker et al., 2013; Fournier et al., 2012). ADMB can fit generic random-effects models (module ADMB-RE) using an EB approach using the Laplace approximation (Skaug and Fournier, 2006). The gradient (i.e., the vector of partial derivatives of the objective function with respect to the parameters) provides a measure of convergence of the parameter estimation procedure in ADMB. By default, ADMB stops when the maximum gradient component is < 10^−4^.

We also tested whether there were noticeable differences in vBGF models when estimating model parameters using a standard non-linear regression fitting routine with no random effects (*nls* function in R) or using ADMB-RE. We carried out this analysis in order to determine whether the fitting of a random-effects model is recommended even when only mean growth trajectories are needed, thus in the case when the fitting of a standard non-linear regression model may represent a theoretically viable procedure.

### 2.4. Population dynamics

We simulated population dynamics of marble, rainbow, and brown trout using individual-based models that include the most important vital rates for the population dynamics for salmonids, i.e. reproduction, juvenile survival (from 0+ to 1+), and survival of fish older than 0+.

Previous studies on the same marble (Vincenzi et al., 2016b), rainbow (Vincenzi et al., 2017a), and brown trout (Vincenzi et al., 2017b) populations have found that that recruitment in all these populations was largely driving variation in population density of fish older than juveniles. Investigations in fish farm have suggested a minimum size for gonads development and reproduction in marble trout (~200 mm) and rainbow trout (~150 mm). However, pedigree reconstruction in four marble trout populations, including LIdri_MT and UIdri_MT (Vincenzi et al., 2017; Vincenzi et al., 2017a), and in the rainbow trout population of Lower Idrijca (LIdri_RT) (Vincenzi et al., 2017a), showed that marble and rainbow trout can reproduce at smaller sizes and reproductive success as number of juveniles produced appears to be independent of parents’ size. Thus, for simulating recruitment (i.e., density of 0+ in September) in the model of population dynamics, we did not use the model of growth and the model of size-dependent fecundity. Instead, we used the coarser-grained stock-recruitment models of Vincenzi et al. (2016b) for marble and rainbow trout, and of Vincenzi et al. (2017b) for brown trout.

Early survival, and in particular the first overwinter survival, is the mayor bottleneck for population size in freshwater salmonids (Vincenzi et al., 2012). Many years of data, and possibly data from multiple populations spanning a wide range of densities (Imre et al., 2005), are necessary to estimate density-dependent survival early in life (from 0+ to 1+). In our model of population dynamics, we used for marble trout the model of density-dependent early survival developed in Vincenzi et al. (2016b). For rainbow and brown trout, we randomly drew at each year of the simulation of population dynamics a value from the set of estimated early survival probabilities reported in Vincenzi et al. (2017a) for rainbow trout and in Vincenzi et al. (2017b) for brown trout.

For modeling survival of fish older than juveniles, we used the population-specific, time-varying survival probabilities estimated in this study. For each population, we simulated 100 years of population dynamics using survival probabilities estimated with final year of sampling *Y*_f_ = 2006, 2008, 2010, 2012, or 2014. At each time step of the simulation of population dynamics, a survival probability was randomly drawn from the *logit* distribution of estimated time-specific survival probabilities and Bernoulli trials were used to determine whether an individual survived or not. Since UVol_BT is a source-sink system (Vincenzi et al., 2017b), we also modeled the influx of brown trout from the source population by doubling the number of fish in each cohort after the first overwinter survival (Vincenzi et al., 2017b).

For each replicate, we recorded (1) mean density of fish older than 0+ over simulation time, and (2) the coefficient of variation (CV) of population density of fish older than 0+ over simulation time. Since freshwater salmonid populations living in Western Slovenia are at contemporary risk of extinction only after the occurrence of extreme climate events such as flash floods or debris flows (Vincenzi et al., 2017; Vincenzi et al., 2017b; Vincenzi et al., 2008), we did not include population extinction as response variable, since the risk of population extinction would almost entirely depend on the modeled intensity and frequency of extreme events (Vincenzi et al., 2008).

For an ensemble of realizations (100 replicates for a fixed set of parameters), we computed: (1) mean and 2.5 and 97.5% quantiles of mean density of fish older than 0+ over simulation time; (2) mean and 2.5 and 97.5% quantiles of CV of density of fish older than 0+ over simulation time.

## 3. Results

Estimates of population densities were variable through time in all four trout populations, with the highest coefficient of variation (CV) for LIdri_RT (0.60) and the lowest for UVol_BT (0.17) (Fig. 1).

For *Y*_f_ > 2006 (i.e., after 6 sampling occasions for marble and rainbow trout, and 5 for brown trout), average survival was constant with varying last year of sampling *Y*_f_ for LIdri_MT, UIdri_MT, and UVol_BT (Fig. 2). For LIdri_RT, average survival was constant for *Y*_f_ > 2009 (i.e., after 12 sampling occasions). The distance between point estimates of maximum and minimum time-varying survival probabilities increased through time, but the 95% CI of maximum and minimum survival probabilities overlapped for each *Y*_f_, except in UVol_BT (Fig. 2).

**Fig. 2.**
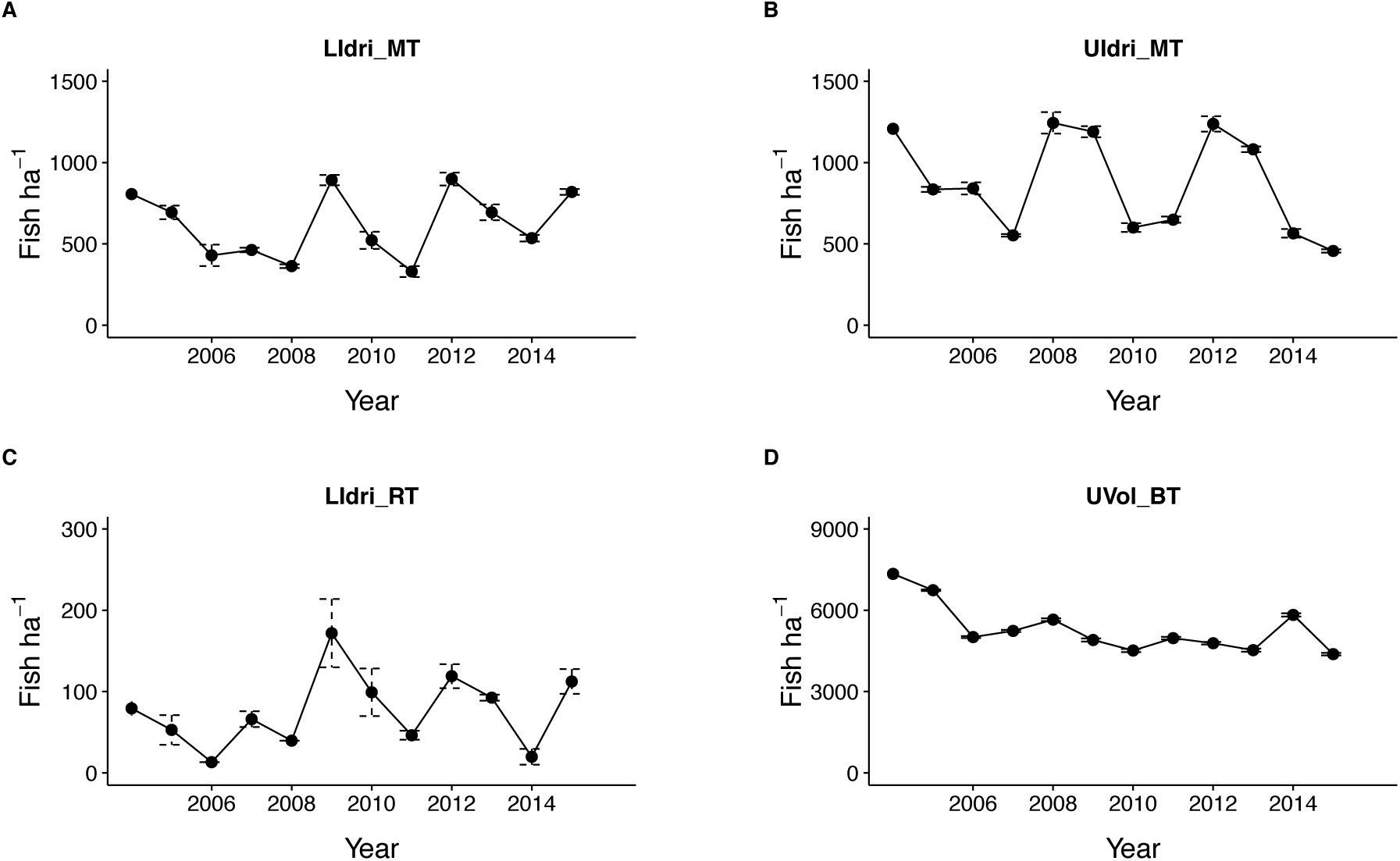
Density over time (individuals older than 0+ in September of each year) of marble trout in Lower (LIdri_MT, panel A) and Upper (UIdri_MT, B) Idrijca, rainbow trout in Lower Idrijca (LIdri_RT, C) and brown trout in Upper Volaja (UVol_BT, D). Dashed vertical lines are 95% confidence intervals. Coefficients of variation of point estimates of population density between 2004 and 2015 were 0.60 (LIdri_RT), 0.17 (UVol_BT), 0.35 (UIdri_MT), and 0.33 (LIdri_RT). Scales on the y-axis are different as estimated densities of rainbow trout and brown trout are much lower and higher, respectively, than those of marble trout.

vBGF models fitted with standard non-linear regression (i.e., without accounting for individual variability in growth) show estimates of asymptotic size that are typically greater than those obtained with random-effects models (Fig. 3). In LIdri_MT, the greater asymptotic size when estimating model parameters without using random-effects was caused by the presence of long-lived individuals that were bigger-at-age than shorter-lived individuals (Fig. 4). In LIdri_RT and UVol_BT, the estimates of asymptotic size when using models with or without random effects were basically the same through time (Fig. 3). The estimates of asymptotic size when using the random-effects vBGF did not change with *Y*_f_, when *Y*_f_ was > 2005 (Fig. 3). For all populations, estimates of parameters in vBGF models with individual random-effects with *Y*_f_ = 2006 or 2014 described the same average growth trajectories (Fig. 5).

**Fig. 3.**
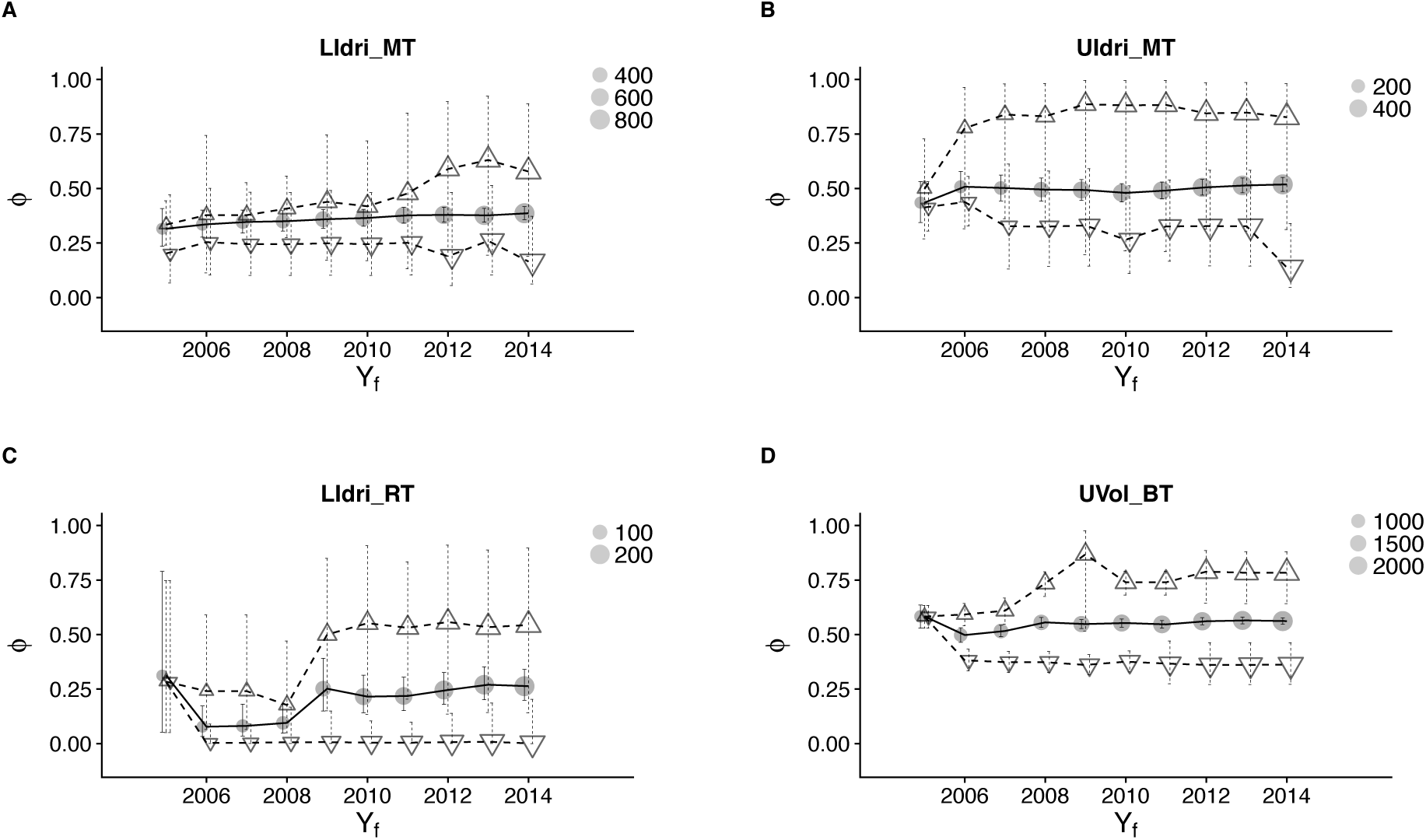
Point estimates ± 95% confidence intervals for average survival (circle), highest (up triangle) and lowest (down triangle) survival for a sampling interval for different last year of sampling *Y*_f_ for LIdri_MT (panel A), UIdri_MT (B), LIdri_RT (C), and UVol_BT (D).

**Fig. 4.**
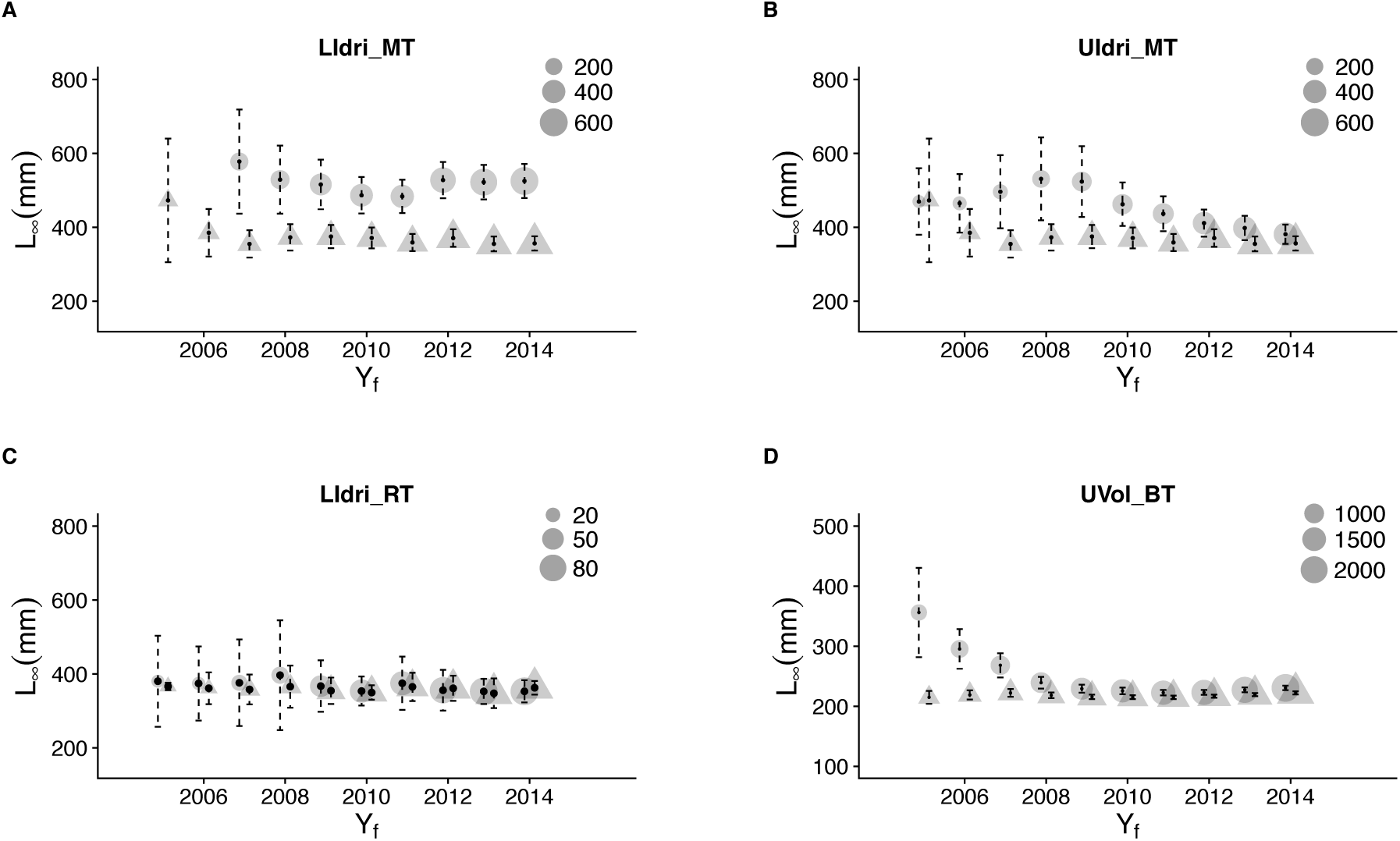
Point estimates ± 95% confidence intervals for asymptotic size *L*_σ_ in the vBGF estimated using standard non-linear regression (circle) or the random-effects model (triangle) for different last year of sampling *Y*_f_ for LIdri_MT (panel A), UIdri_MT (B), LIdri_RT (C), and UVol_BT (D). Size of symbols is proportional to the number of unique individuals in the data set. For LIdri_MT, the estimate of asymptotic size when using standard non-linear regression techniques with *Y*_f_ = 2005 and 2006 (not shown) were (mean[95%CI]): 1818 mm [(−1899)−5535] and 1555 mm [(−577)−3688], respectively.

**Fig. 5.**
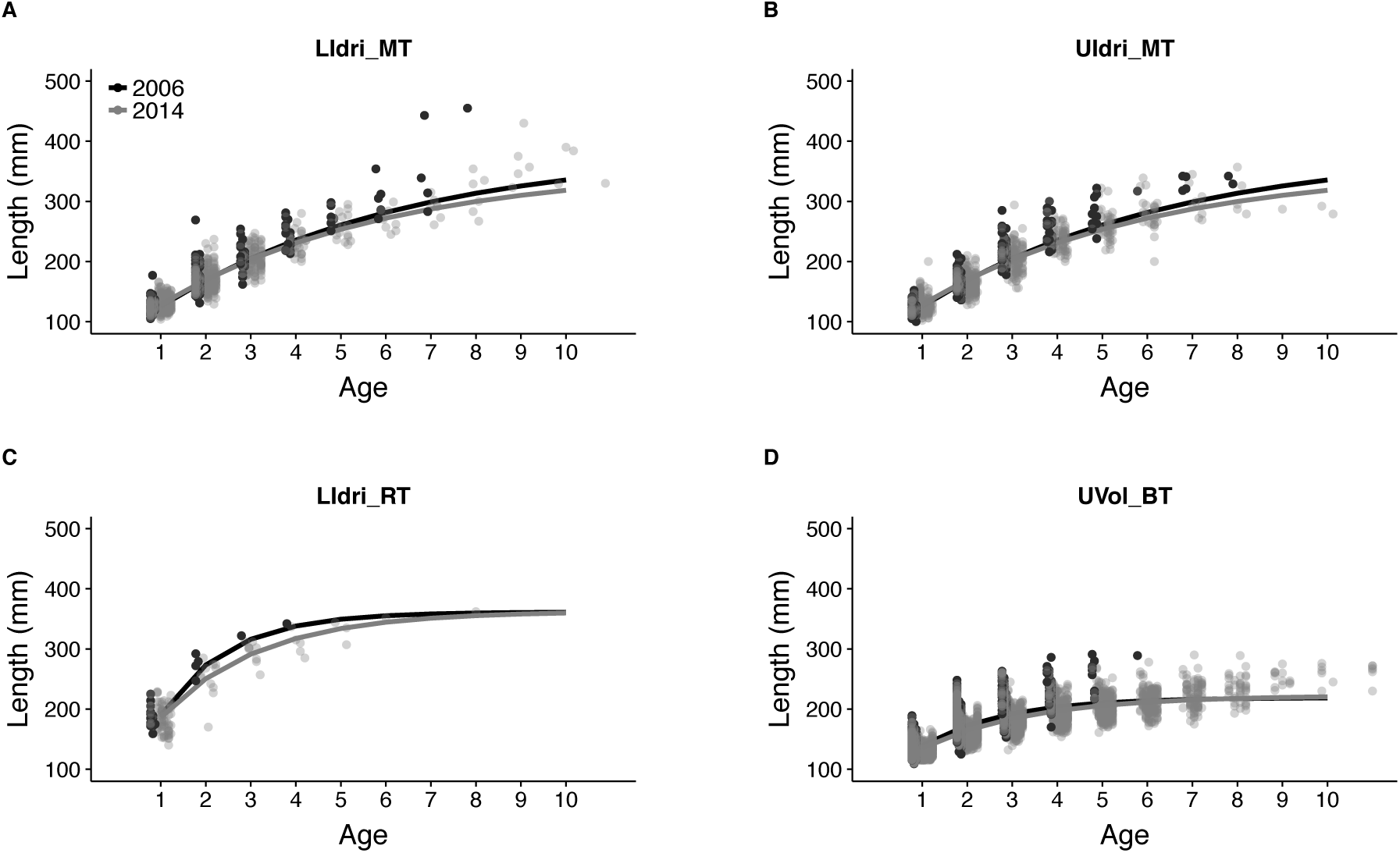
Average growth trajectories estimated using the random-effects vBGF models using data collected up to 2006 (black) and up to 2014 (gray) for LIdri_MT (panel A), UIdri_MT (B), LIdri_RT (C), and UVol_BT (D).

**Fig. 6.**
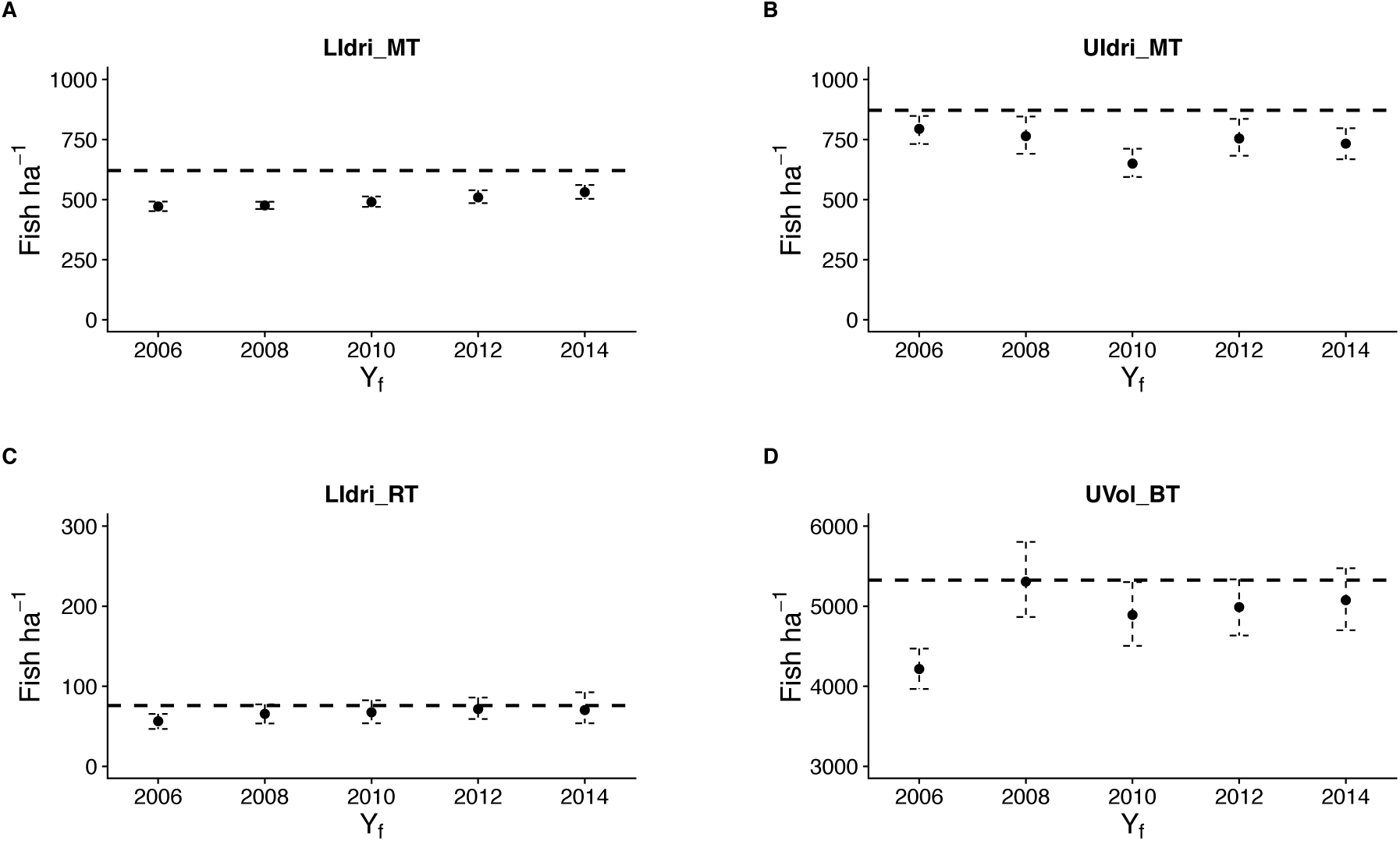
Observed mean population density of fish older than 0+ from 2004 to 2015 (horizontal dashed line) and mean and 2.5 and 97.5% quantiles of mean density of fish older than 0+ over simulation time for different last year of sampling *Y*_f_ for LIdri_MT (panel A), UIdri_MT (B), LIdri_RT (C), and UVol_BT (D). Scales on the y-axis are different as estimated densities of rainbow trout and brown trout are much lower and higher, respectively, than those of marble trout.

**Fig. 7.**
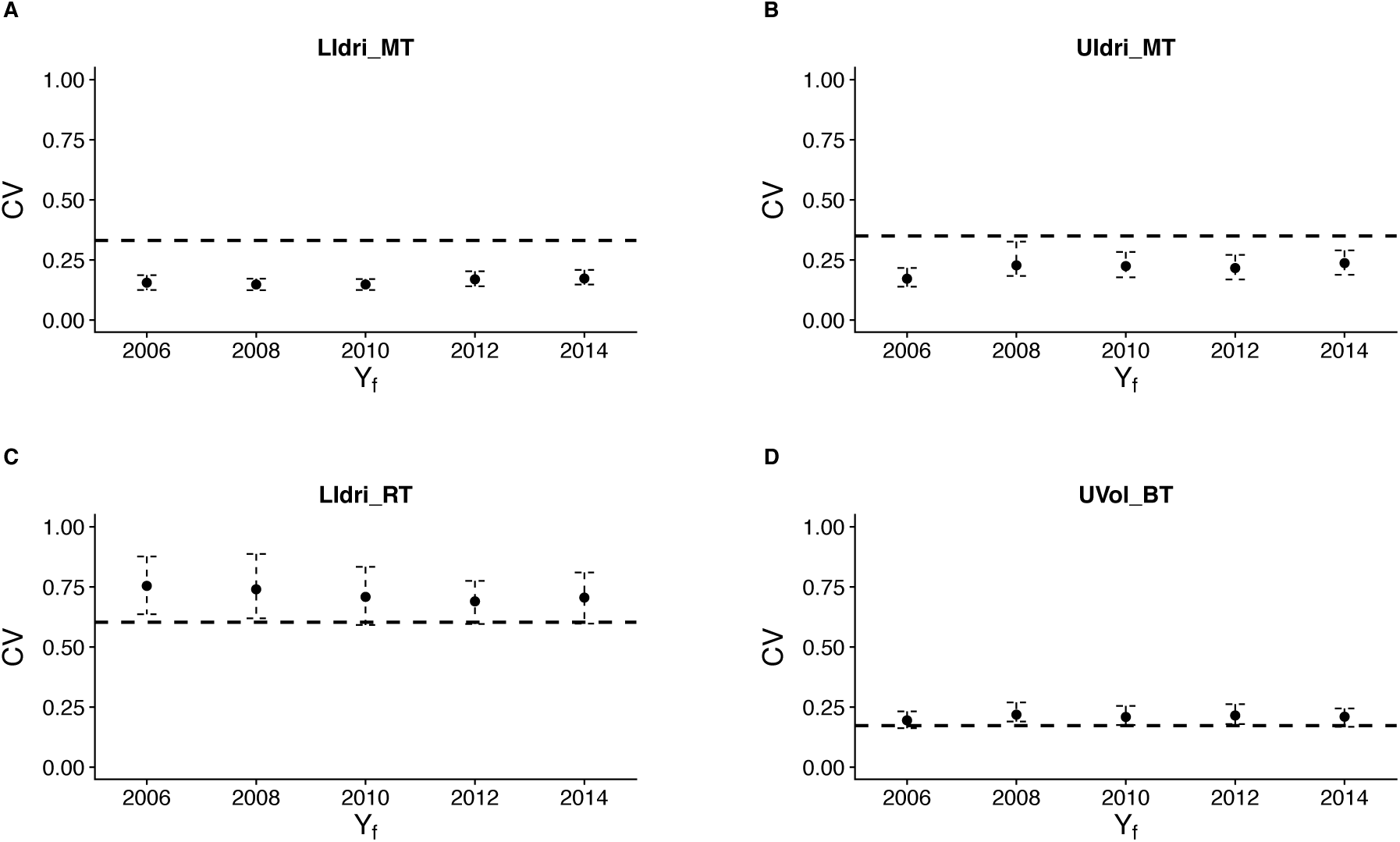
Observed coefficient of variation of density of fish older than 0+ from 2004 to 2015 (horizontal dashed line) and mean and 2.5 and 97.5% quantiles of coefficient of variation of density of fish older than 0+ over simulation time for different last year of sampling *Y*_f_ for LIdri_MT (panel A), UIdri_MT (B), LIdri_RT (C), and UVol_BT (D).

Using time-varying fish survival probabilities estimated at different *Y*_f_ had little effects on predictions of mean population density and its coefficient of variation. The simulated mean densities were within 10% of the observed mean densities for all trout populations for all *Y*_f_. The only exception was UVol_BT, for which the simulated mean densities with *Y*_f_ = 2006 were 20% lower than the observed mean density over 2004-2015. CV of density from simulations was lower than the observed CV over 2004-2015 for LIdri_MT, and similar to the observed CV for the other trout populations.

## 4. Discussion

Effective conservation of species requires the estimation of variation in vital rates and life-history traits, an understanding of the determinants of the observed variation, and the integration of vital rates and life histories estimated from focal data, findings from published literature, and controlled experiments into models of population dynamics for prediction and evaluation of scenarios. Using data from, loosely defined, short-or long-term monitoring may lead to models with parameter estimates of varying accuracy due to the availability and quality of data sets (Beissinger and Westphal, 1998). Thus, modeling of population dynamics is a valuable tool for understanding ecological systems and directing conservation strategies, but how long monitoring programs informing those models should go on is often unclear.

In this work, we investigated how estimates of vital rates and predictions of population dynamics change along the monitoring program with the collection of more data. We found that in 3 out of 4 populations, a few years of data (3 or 4 sampling occasions, close to the generation time of marble trout and ~ 1.5 times the generation time of rainbow and brown trout) provided the same estimates of average survival and growth as those obtained by more than 15 sampling occasions, while the estimation of the range of survival probabilities (i.e., ideally the distribution of survival probabilities through time, more often the distance between maximum and minimum survival) required, as expected, more sampling occasions (up to 22 for marble trout), with little reduction of uncertainty around the point estimates. Predictions of mean density and variation in density over time did not change with more data after the first 5 years (i.e., 10 sampling occasions) and were within 10% of the observed mean and variation in density over 11 years.

### 4.1. Survival

Previous work has found cohort and time effects on the survival probabilities of freshwater salmonids living in Western Slovenian streams (Vincenzi et al., 2017a, 2017b; Vincenzi et al., 2016b). As it was found that neither water temperature nor population density seemed to explain variation in survival, the observed variation might be ascribed to variation in flow rates, trophic conditions, or other unobserved/unmeasured properties of the environment (Vincenzi et al., 2016b). Although due to the effects of sample size and of a fairly stable environment in Lower Idrijca, Upper Idrijca, and Upper Volaja we expected the marginal effect of additional data to be increasingly smaller along the monitoring program, we found that even after 6 to 8 sampling occasions the estimates of average survival (both point estimates and confidence intervals) did not change with additional years of data. The only exception was the rainbow trout population of Lower Idrijca - the smallest of the four salmonid populations -, for which the estimates of average survival remained stable over time only after 12 sampling occasions. Overall, capture-recapture data from between 300 and 500 tagged fish were sufficient to have stable estimates of average survival probabilities. Since newly tagged fish entered the data set at each sampling occasion, further studies will investigate how many complete life histories are needed to obtain stable estimates of average survival probabilities.

Due to small population sizes, the confidence intervals of the estimates of maximum and minimum survival over a sampling interval overlapped in all populations except the brown trout population of Upper Volaja. In addition, while maximum survival is expected to be have a ceiling determined by the habitat conditions and the ecology of the species that can be estimated with a few years of data, the estimation of very low survival probabilities such as those caused by flash floods and debris flows (Vincenzi et al., 2017; Vincenzi et al., 2016b) may require decades-long monitoring programs.

### 4.2. Growth

Estimates of body growth are fundamental for management. For instance, age-structured stock assessment methods are based on sizes-at-age that are often derived from parameters of the von Bertalanffy growth model for that species (Katsanevakis and Maravelias, 2008). Size-at-age, which is the easiest-to-observe realization of the growth process, often varies considerably among individuals living in the same environment. In the four trout populations, the size of the smallest age-1 fish was ~50% of the size of the biggest age-1 fish. This variation may result from complex interactions between genetic, environmental, population factors, and chance. When not accounted for, the presence of individual variation can bias the estimation of growth and other demographic traits (Vincenzi et al., 2016a, 2014).

Longitudinal data (e.g., tag-recapture) and random-effects models greatly facilitate the estimation of individual and group (i.e., sex, year-of-birth) variation in growth. In particular for the two marble trout populations, we found that standard non-linear regression (i.e., models without random-effects) provided estimates of asymptotic size that were consistently larger (up to *Y*_f_ = 2014 for marble trout in Lower Idrijca and up *Y*_f_ = 2012, ~ 800 tagged fish, for marble trout in Upper Idrijca) than those provided by the random-effects vBGF models. In marble trout, the main type of intra-specific competition for resources seems to be interference competition for space (Vincenzi et al., 2016a), probably due to high territoriality. In interference competition, bigger individuals (in the case of marble trout, those with access to better sites) reduce the access to resources, such as space and food, of smaller individuals, and may also live longer than smaller individuals. On the other hand, the estimates of asymptotic size using the random-effects vBGF were little affected by the use of more data when 100 to 200 individuals of various ages where already included in the data set. Since the vBGF parameters can seldom be interpreted separately (Vincenzi et al., 2014), the analysis of the whole growth trajectories is crucial for understanding variation in growth. Also when examining whole growth trajectories, we found that the growth trajectories predicted by the random-effects vBGF models were basically identical when using data from 3 (100 to 200 individuals of various ages) or 11 years of monitoring.

### 4.3. Population dynamics

In our model of population dynamics, the differences between predictions of mean density and its variation when using data from 3 (6 sampling occasions) or 11 (22 sampling occasions) were negligible in all populations except the brown trout population of Upper Volaja. The low mean population density predicted for Upper Volaja when using data from 5 sampling occasions was due to low probabilities of survival in Upper Volaja between 2004 and 2007, whose inclusion in the model of population dynamics led to predictions of mean population size much lower than those provided by models parameterized with more data and more representative environmental conditions. Thus, when the estimates of survival probabilities are not representative of the conditions typically experienced by individuals, the predictions of the model can be largely inaccurate, as in the case of the brown trout population of Upper Volaja.

In our analyses, only survival probabilities were different among models. However, the parameters with the most available data are not necessarily the parameters that have the greatest effect on model predictions. In some cases, empirical data may be lacking for parameters that can substantially alter model predictions. In our study, we were able to include a model of density-dependent early survival only in the models of population dynamics for the two populations of marble trout, since even more than 10 years of data were not sufficient to estimate parameters of similar models for brown trout and rainbow trout. However, randomly drawing an early survival probability at each time step from the set of estimated probabilities did not compromise the accuracy of the predictions of mean population density.

As for recruitment dynamics, the models in Vincenzi et al. (2016b) for marble and rainbow trout and in Vincenzi et al. (2017b) for brown were able to explain only part of the variability in recruitment (i.e., < 30%). The relative balance between spawning stock size and environmental factors as determining recruitment in freshwater salmonids is still debated and probably context-specific (Einum, 2005; Nicola et al., 2008). Recruitment in marble, rainbow, and brown trout was highly variable over time; the marble and rainbow trout populations of Lower and Upper Idrijca were recruitment-driven, as indicated by the strong one-year lagged correlation between density of older than newborn trout and density of newborns (Vincenzi et al., 2017a; Vincenzi et al., 2016b). In Upper Volaja, the absence of a correlation between density of older than newborn trout and density of newborns was caused by immigration of 0+ and 1+ from the source population (Vincenzi et al., 2017b). Despite the uncertainty in recruitment models, the predictions of mean population density and variation of density over time were largely accurate.

## Acknowledgements

We thank the employees and members of the Tolmin Angling Association (Slovenia) for carrying out fieldwork since 1993.

## Data Statement

Data and R code: https://github.com/simonevincenzi/Limit_sampling

